# Utilizing artificial intelligence system to build the digital structural proteome of reef-building corals

**DOI:** 10.1101/2022.06.27.497859

**Authors:** Yunchi Zhu, Xin Liao, Tingyu Han, J.-Y. Chen, Chunpeng He, Zuhong Lu

## Abstract

Reef-building corals play an important role in the marine ecosystem, and analyzing their proteomes from a structural perspective will exert positive effects on exploring their biology. Here we integrated mass spectrometry with newly published ColabFold to obtain digital structural proteomes of dominant reef-building corals. 8,382 proteins co-expressed in *A. muricata, M. foliosa* and *P. verrucosa* were identified, then 8,166 of them got predicted structures after around 4,060 GPU hours of computation. The resulting dataset covers 83.6% of residues with a confident prediction, while 25.9% have very high confidence. Our work provides insight-worthy predictions for coral research, confirms the reliability of ColabFold in practice, and is expected to be a reference case in the impending high-throughput era of structural proteomics.

## Introduction

Coral reefs serve as a living environment for more than 30% of marine animals and plants [1–2], while they are suffering from sharply declining due to global warming, changes in the physicochemical environment of the ocean, and massive encroachment of the predatory crown-of-thorns starfish [3–7]. Several researchers even propose that features similar to those exhibited during the last mass extinction have emerged in scleractinian coral populations, including population shrinkage, transplanting of colonies to the aphotic zone, and zygote dormancy [8–9]. Such serious situation has brought about research focused on the growth, restoration, and ecological defence of reef-building corals.

However, in contrast to more common model organisms, there remains a lack of public omics data from reef-building corals, no exception for proteome bridging physiological function and genome. Taking the UniProt database as an example, as of June 2022, it has collected no more than 50 reviewed proteins from *Acropora*, one of the most species rich coral genera, while for another two dominant genera *Montipora* and *Pocillopora*, the number even fails to reach 5. Various factors such as geographic location and technical limitations result in the “data gap” for coral research [10], and to make matters worse, COVID-19 has exacerbated the risks of sampling at wild as well as performing “wet” experiments. Mass spectrometry technology having been applied in coral proteomics research [11–14] enables researchers to obtain high-throughput protein information from relatively small samples in standardized steps, yet according to traditional protocols downstream analysis on protein structures and functions still requires cumbersome manual operations.

The booming AI (artificial intelligence) technology is expected to provide new solutions for current predicament. The results of the biennial Critical Assessment of protein Structure Prediction (CASP) have revealed the substantial progress in protein structure prediction [15] and brought AlphaFold2 [16] to spotlight. This AI system achieved a record score of 92.4 in CASP14 (2020), beating all competing models. As the structure modelling solution challenging x-ray crystallography and cryo-electron microscopy, AlphaFold2 can directly transform sequences to structures with high accuracy, particularly beneficial for studies on several non-model organisms represented by corals. These species may be important to our ecology and society, but existing experimental protocols for them are not as perfect as those for model organisms, moreover people tend to focus on their potentially valuable components rather than a head-to-tail understanding of them. Assisted by AI modelling, scientists can rapidly acquire their structural proteomes [17] in the digital lab, then use computational biology methods to find key proteins and explore crucial physiological functions, so as to pave the way for applications including breeding and protection. Such “dry-led” strategy might not only reduce experimental costs but also better protect researchers’ health during COVID-19.

Compared to other prediction algorithms such as RoseTTAFold [18], AlphaFold2 has obvious disadvantages in performance, even domains could consume long computation time [19]. Fortunately, its open source has attracted joint efforts of developers to improve it. In 2021, Zhong and colleagues from the center for high performance computing (HPC) of Shanghai Jiao Tong University released ParaFold [20], the specific AlphaFold version for their HPC clusters, making a successful attempt to accelerate AlphaFold2. In June 2022, ColabFold [21] claiming to make protein folding accessible to all got officially published. Replacing Jackhmmer with MMseqs2 [22] and utilizing optimized model, ColabFold can run dozens of times faster than original AlphaFold2, which means it greatly expands the throughput of protein structure modelling, thus breaking the last hurdle in effectively predicting complete structural proteome.

Here we integrated mass spectrometry with AI system to obtain digital structural proteomes of dominant reef-building corals. Deploying the latest release of ColabFold on the Big Data Computing Center of Southeast University, we predicted structures of more than 8,000 co-expressed proteins among *A. muricata, M. foliosa* and *P. verrucosa* in approximately 4,060 GPU hours. The resulting dataset named CP-8382 covers 83.6% of residues with a confident prediction and 25.9% with very high confidence. We also developed a search engine interface in the style of AlphaFold Protein Structure Database [23] to open our data to the community (http://corals.bmeonline.cn/prot/).

## Materials and Methods

### Experimental model and subject details

The species including *A. muricata, M. foliosa* and *P. verrucosa* in the study were collected from the Xisha Islands in the South China Sea (latitude 15°40′–17°10′ north, longitude 111°–113° east).

The coral samples were cultured in our laboratory coral tank with conditions conforming to their habitat environment. All the species were raised in a RedSea^®^ tank (redsea575, Red Sea Aquatics Ltd) at 26°C and 1.025 salinity (Red Sea Aquatics Ltd). The physical conditions of the coral culture system are as follows: three coral lamps (AI^®^, Red Sea Aquatics Ltd), a protein skimmer (regal250s, Reef Octopus), a water chiller (tk1000, TECO Ltd), two wave devices (VorTech™ MP40, EcoTech Marine Ltd), and a calcium reactor (Calreact 200, Reef Octopus), etc.

### Total Protein Extraction

Sample was ground individually in liquid nitrogen and lysed with PASP lysis buffer (100 mM NH_4_HCO_3_, 8 M Urea, pH 8), followed by 5 min of ultrasonication on ice. The lysate was centrifuged at 12000 g for 15 min at 4°C and the supernatant was reduced with 10 mM DTT for 1h at 56°C, and subsequently alkylated with sufficient IAM for 1 h at room temperature in the dark. Then samples were completely mixed with 4 times volume of precooled acetone by vortexing and incubated at −20°C for at least 2h. Samples were then centrifuged at 12000 g for 15 min at 4°C and the precipitation was collected. After washing with 1mL cold acetone, the pellet was dissolved by dissolution buffer (8 M Urea, 100 mM TEAB, pH 8.5).

### Protein Quality Test

BSA standard protein solution was prepared according to the instructions of Bradford protein quantitative kit, with gradient concentration ranged from 0 to 0.5 g/L. BSA standard protein solutions and sample solutions with different dilution multiples were added into 96-well plate to fill up the volume to 20 μL, respectively. Each gradient was repeated three times. The plate was added 180 μL G250 dye solution quickly and placed at room temperature for 5 minutes, the absorbance at 595 nm was detected. The standard curve was drawn with the absorbance of standard protein solution and the protein concentration of the sample was calculated. 20 μg of the protein sample was loaded to 12% SDS-PAGE gel electrophoresis, wherein the concentrated gel was performed at 80 V for 20 min, and the separation gel was performed at 120 V for 90 min. The gel was stained by coomassie brilliant blue R-250 and decolored until the bands were visualized clearly.

### TMT Labeling of Peptides

Each protein sample was taken and the volume was made up to 100 μL with DB dissolution buffer (8 M Urea, 100 mM TEAB, pH 8.5). Trypsin and 100 mM TEAB buffer were added, sample was mixed and digested at 37 °C for 4h. And then, trypsin and CaCl_2_ were added, sample was digested overnight. Formic acid was mixed with digested sample, adjusted pH under 3, and centrifuged at 12000 g for 5 min at room temperature. The supernatant was slowly loaded to the C18 desalting column, washed with washing buffer (0.1% formic acid, 3% acetonitrile) 3 times, then eluted by some elution buffer (0.1% formic acid, 70% acetonitrile). The eluents of each sample were collected and lyophilized. 100 μL of 0.1 M TEAB buffer was added to reconstitute, and 41 μL of acetonitrile-dissolved TMT labeling reagent was added, sample was mixed with shaking for 2 h at room temperature. Then, the reaction was stopped by adding 8% ammonia. All labeling samples were mixed with equal volume, desalted and lyophilized.

### Separation of fractions

Mobile phase A (2% acetonitrile, adjusted pH to 10.0 using ammonium hydroxide) and B (98% acetonitrile) were used to develop a gradient elution. The lyophilized powder was dissolved in solution A and centrifuged at 12,000 g for 10 min at room temperature. The sample was fractionated using a C18 column (Waters BEH C18, 4.6×250 mm, 5 μm) on a Rigol L3000 HPLC system, the column oven was set as 45°C. The detail of elution gradient was shown in **Table S1**. The eluates were monitored at UV 214 nm, collected for a tube per minute and combined into 10 fractions finally. All fractions were dried under vacuum, and then, reconstituted in 0.1% (v/v) formic acid (FA) in water.

### LC-MS/MS analysis

For transition library construction, shotgun proteomics analyses were performed using an EASY-nLC™ 1200 UHPLC system (Thermo Fisher) coupled with a Q Exactive™ series mass spectrometer (Thermo Fisher) operating in the data-dependent acquisition (DDA) mode. 1 μg sample was injected into a home-made C18 Nano-Trap column (4.5 cm×75 μm, 3 μm). Peptides were separated in a home-made analytical column (15 cm×150 μm, 1.9 μm), using a linear gradient elution as listed in **Table S2**. The separated peptides were analyzed by Q Exactive™ series mass spectrometer (Thermo Fisher), with ion source of Nanospray Flex™ (ESI), spray voltage of 2.3 kV and ion transport capillary temperature of 320°C. Full scan ranges from m/z 350 to 1500 with resolution of 60000 (at m/z 200), an automatic gain control (AGC) target value was 3×10^6^ and a maximum ion injection time was 20 ms. The top 40 precursors of the highest abundant in the full scan were selected and fragmented by higher energy collisional dissociation (HCD) and analyzed in MS/MS, where resolution was 45000 (at m/z 200) for 10 plex, the automatic gain control (AGC) target value was 5×10^4^ the maximum ion injection time was 86 ms, a normalized collision energy was set as 32%, an intensity threshold was 1.2×10^5^, and the dynamic exclusion parameter was 20 s.

### Protein identification and quantitation

The resulting spectra from each run were searched separately against protein-coding sequences from PRJNA544778 by Proteome Discoverer 2.2 (PD 2.2, Thermo) [24]. The searched parameters are set as follows: mass tolerance for precursor ion was 10 ppm and mass tolerance for product ion was 0.02 Da. Carbamidomethyl was specified as fixed modifications, Oxidation of methionine (M) and TMT plex were specified as dynamic modification, acetylation and TMT plex were specified as N-Terminal modification in PD 2.2. A maximum of 2 miscleavage sites were allowed.

In order to improve the quality of analysis results, the software PD 2.2 further filtered the retrieval results: Peptide Spectrum Matches (PSMs) with a credibility of more than 99% were identified PSMs. The identified protein contains at least 1 unique peptide. The identified PSMs and protein were retained and performed with FDR no more than 1.0%. The protein quantitation results were statistically analyzed by T-test.

### Structure modelling

Protein sequences were sorted by length and numbered as CPXXXXXXXX according to order before sent to ColabFold (1.3.0) platform. 1,053 proteins no longer than 200 aa were calculated on NVIDIA Tesla P100 while others were calculated on NVIDIA Tesla V100 cluster of the Big Data Computing Center of Southeast University. The parameters of ColabFold were set to --*amber*, --*templates*, --*num-recycle 3*, --*use-gpu-relax*. For each protein, structure with highest pLDDT scores (*\*_relaxed_rank_1_model_x.pdb*) was preserved and labeled as CPXXXXXXXX.pdb.

### Search engine development

Elasticsearch 7.12.1 was employed as the key module. Each coral protein was annotated by NR (diamond v2.0.14.152, *blastp --evalue 1e-5 -k 1*), UniProt (diamond v2.0.14.152, *blastp* -- *evalue 1e-5 -k 1*) and InterPro (interproscan-5.54-87.0), then annotations were transformed into keywords in Elasticsearch index. Web interface was implemented using PHP 7.2, while Nginx worked as web server.

Sequenceserver [25] 2.0.0 were deployed as the BLAST server, source codes of which were modified to link each hit to its information page. Mol* [26] in the style of AlphaFold Protein Structure Database was imported as structure viewer.

## Results and Discussion

### Protein identification and annotation

8,382 proteins co-expressed in *A. muricata, M. foliosa* and *P. verrucosa* were identified by Proteome Discoverer, sequences and expression profile of which are shown in **Table S3**. It is assumed that a considerable portion of proteomes be conserved among these dominant coral genera, for the total protein-coding gene number of one scleractinian coral is unlikely to exceed 25,000 according to previous reports [9–10][27]. Over 97% of them are shorter than 2,600 aa, while the longest is up to 14,622 aa.

Proteins were annotated with NR and UniProt for homologs, and InterPro for domains. As illustrated in **Fig S1.A**, NR annotations map most proteins to scleractinians, not other cnidarians or marine organisms, indicating the increasing efforts into filling the “data gap” hindering coral research [10] in recent years. **Fig S1.B** shows some domains frequently found on coral proteins. EF-hand domain related to calcium signalling pathways [28] gets the top 1, just consistent with these corals’ roles as reef builders, who deal with calcium ion every day for skeleton construction and homeostatic regulation. All annotations were employed as keywords for searching.

### Protein structure prediction

After about 4,060 GPU hours of computation, ColabFold succeeded to generate 8,166 structures, covering 97.4% of our coral protein dataset and touching the maximum length that an NVIDIA Tesla V100 (32 GB) can process. The model confidence distribution is presented in **Fig 1**. In resulting dataset 83.6% of residues have pLDDT (predicted local distance difference test) larger than 70, which are considered as confident predictions [15], and 25.9% get very high pLDDT over 90. No more than 2% of predicted structures have an average pLDDT below 50. The predictions from ColabFold can be recognized as credible overall.

**Fig 1.**
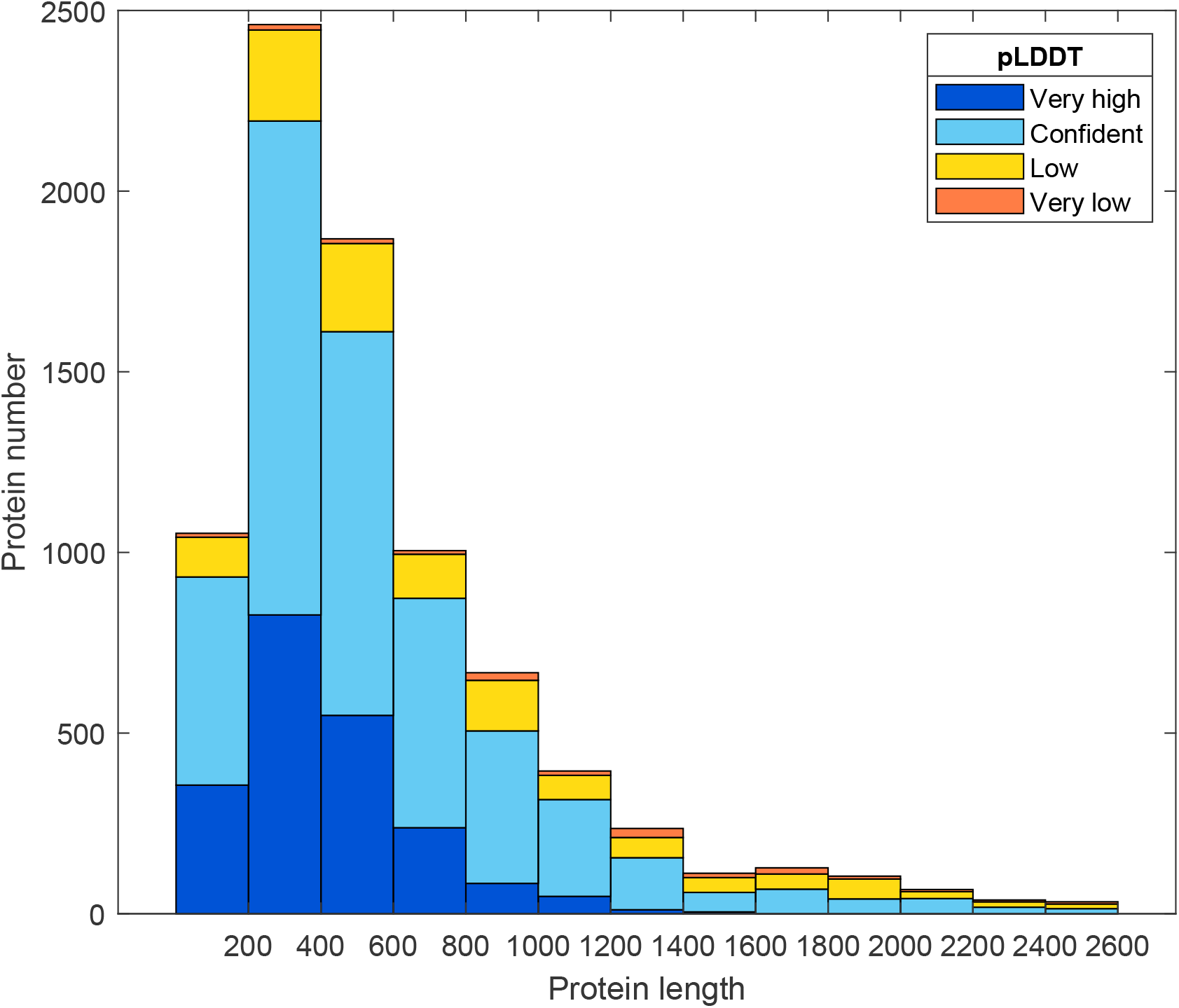
Distribution of model confidence against protein length. Horizontal axis is protein length and vertical axis is protein number. Model confidence calculated by pLDDT are color-coded. Very high: pLDDT > 90; Confident: 90 > pLDDT > 70; Low: 70 > pLDDT > 50; Very low: pLDDT < 50.

**Fig 2** demonstrates several highlighted structure predictions from the resulting dataset. There are few doubts that biomineralization is the most significant function of reef-building corals [29–30], thus their skeletal proteome regulating the mineral deposition is always of interest to scientists. Our results cover most of recognized coral mineralization-related proteins, including skeletal organic matrix protein (SOMP), skeletal aspartic acid-rich protein, collagen, carbonic anhydrase, etc. [11–12] **Fig 2.A** shows the sequence and structural alignment among three acidic SOMPs, two identified in this work and one from UniProt (B3EWY7). Although the sequence identity is only 40%-60%, it appears that their structures are broadly similar, consisting of an Asp-rich tongue-like region and a β-sheet-formed region. According to previous studies, Asp-rich proteins are supposed to interact directly with calcium carbonate crystals promoting crystal nucleation, determining the growth axes and inhibiting the crystal growth [31–32], and they usually have high-capacity yet low-affinity calcium-binding properties [33]. Our structure predictions may provide an explanation that for these proteins Asps are concentrated in an open tongue-like region, reducing the steric hindrance while weakening the binding strength with calcium ions. Besides, the β-sheet-rich region at the base of the “Asp tongue” form a barrel-like local structure, which is also found in many other uncharacterized skeletal organic matrix proteins (USOMP) (**Fig S2**). These barrel-like regions probably have transmembrane functions [34], however existing methods fail to annotate them with any known domains, making these SOMPs uncharacterized. This phenomenon might not only urge biologists to gain deeper insights into protein domains, but also enlighten bioinformaticians to design novel annotation algorithms for 3D structure data.

**Fig 2.**
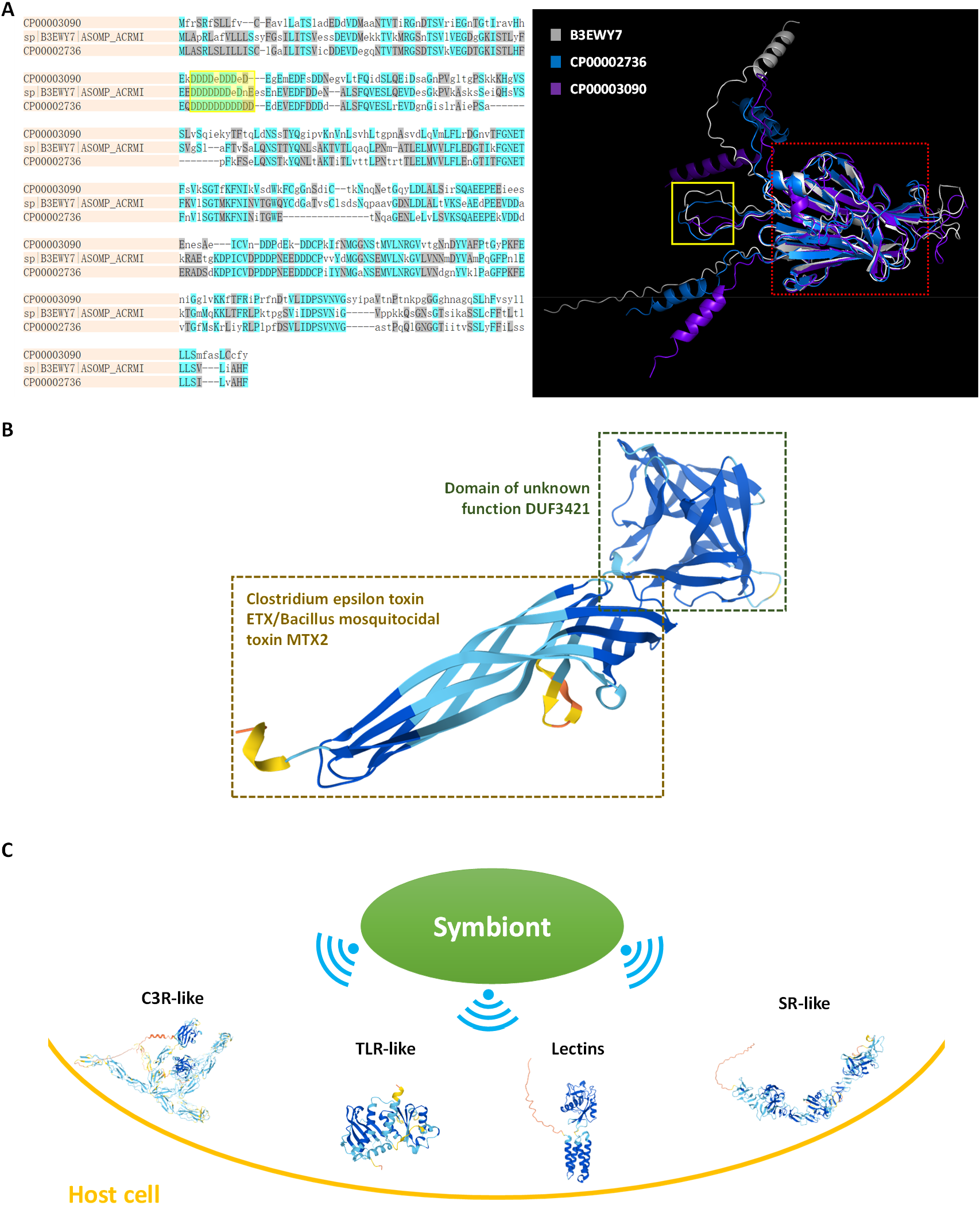
Highlighted structure predictions. **A)** Sequence and structure alignment among CP00002736, CP00003090 and B3EWY7. Asp-rich regions were framed in yellow and β-sheet-formed regions were framed in red. **B)** Structure of CP00002607 similar to natterin-4. **C)** Structures of several pattern recognition receptors involved in host-symbiont signalling, including C3R (complement 3 receptor), TLR (toll-like receptor), lectin and SR (scavenger receptor).

Toxins are widely employed by marine organisms for prey capture and defense of the territory [35]. Natterin proteins discovered in the venom of the medically significant Brazilian toadfish *Thalassophryne nattereri* [36] are representative perforin-like toxins which can insert into the lipid bilayer to trigger the disruption of membrane function [37]. We identify one co-expressed Natterin-4-like protein CP00002607, as illustrated in **Fig 2.B**. The drill-like toxic domain is supposed to “bore holes” on membranes, resulting in electrolyte leakage and inflammation [38]. The function of DUF3421 has not been commonly recognized [39], though several researchers propose such DM9-containing proteins be new members of PRRs (pattern recognition receptors) [40–42]. From a structural point of view, this DM9-containing domain is at the end of “toxic drill” with a funnel-like cavity. It indeed has potential to bind signalling molecule playing roles in the antagonism between toxin and immune system, nevertheless the possibility that it just exacerbates membrane damage or attaches other toxic chemicals cannot be ruled out. This structure prediction might help to deepen our understanding of marine biotoxin and provide inspiration for drug development [43], not to mention that it can serve as raw data for molecular docking itself.

Symbiosis is another important topic for coral biologists and ecologists, as reef-building corals obtain the majority of their energy and nutrients from their algal symbionts (mainly Symbiodiniaceae), and loss of symbionts is causing coral bleaching threatening human survival. It is recognized that improved knowledge of interpartner signalling in coral holobionts could be applied to solutions against the coral reef crisis [44]. Previous studies reveal that the initiation of coral symbiosis depends on the interaction between PRRs on host gastrodermal nutritive phagocytes and MAMPs (microbe-associated molecular patterns) of Symbiont [45–46], as shown in **Fig 2.C**. Complete structures of several representative PRRs, such as C3R (complement 3 receptor), TLR (toll-like receptor), lectin and SR (scavenger receptor), get confident prediction in our work with pLDDT ranging from 75 to 88. Unfortunately, our predictions fail to cover PRRs too large for GPU devices to process, and symbiont proteins including MAMPs are beyond the scope of our experiment design. It will be an excellent work to acquire Symbiodiniaceae structural proteomes and integrate them with corals’, which may bring about novel insights into coral symbiosis and interpartner signalling in cnidarians. In fact, the “data gap” for Symbiodiniaceae turns out to be even larger than that for corals [10], presenting both challenges and opportunities.

### Web interface of CP-8382

Combining annotations and predicted structures, the resulting dataset was named as CP-8382 and cruated into an online database (http://corals.bmeonline.cn/prot/). **Fig 3** describes its data content as well as the workflow of corresponding web interface. Predicted structures are displayed in the style of AlphaFold Protein Structure Database, where pLDDT of each residue is marked by color, and Mol* app enables users to zoom, rorate or take screenshot. They can be downloaded in the format of PDB or CIF. PAE (Predicted aligned error) graphs are given for assessing confidence in global features [15], raw data of which can be accessed in JSON format. Annotations with keywords highlighted are also presented. All the above information is accessible via search engine or BLAST server at the web interface. **Video S1** provides a demo of searching skeletal aspartic acid-rich protein in CP-8382.

**Fig 3.**
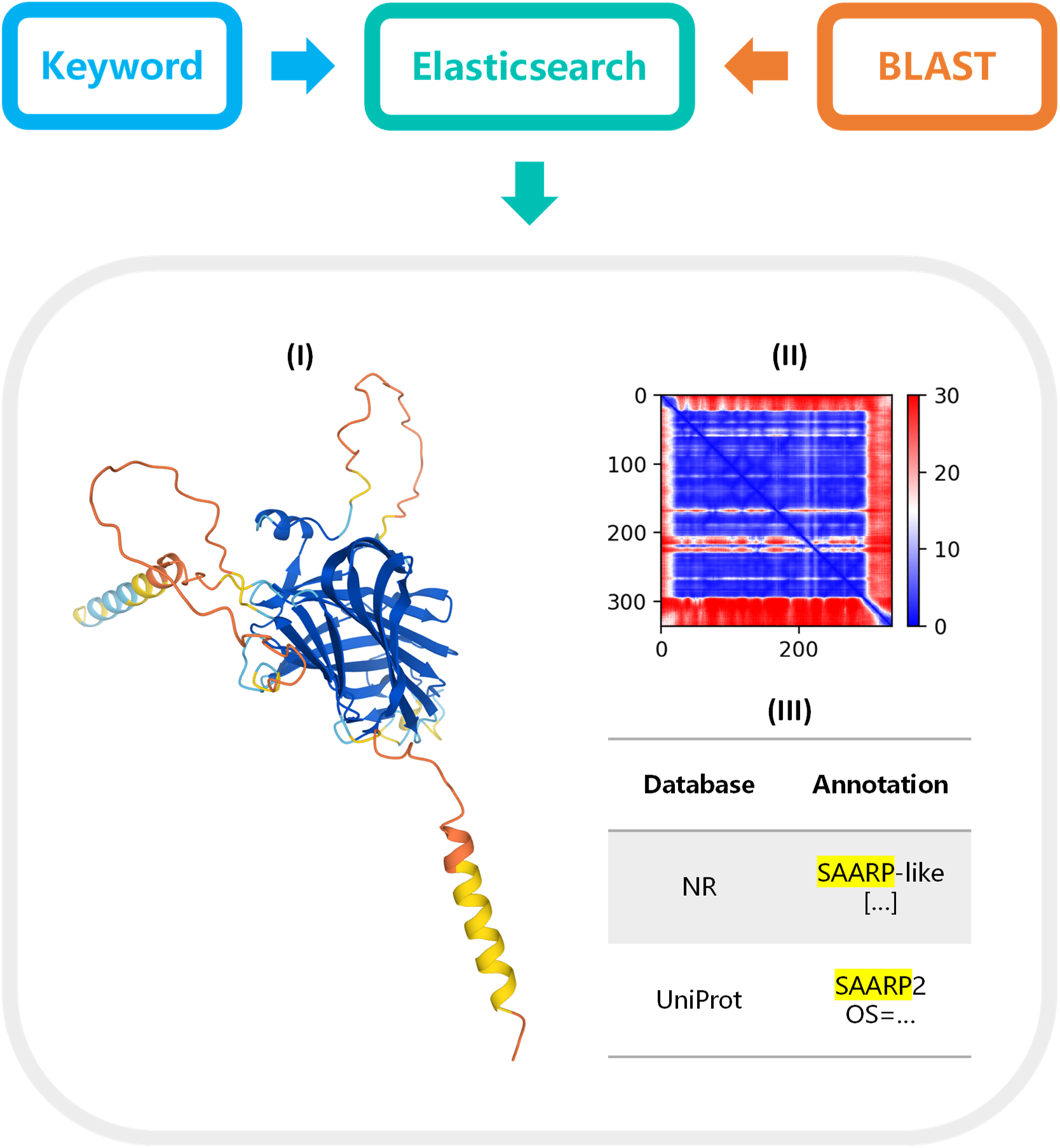
Content of CP-8382 dataset and workflow of its web interface. Users can utilize search engine or BLAST server to acquire their interested coral proteins on the website, where elasticsearch module will process inputs and return the following information: (I) Predicted structure displayed in the style of AlphaFold Protein Structure Database; (II) PAE graph, where the colour at position (x, y) indicates AlphaFold’s expected position error at residue x when the predicted and true structures are aligned on residue y; (III) Annotations with keywords highlighted.

## Conclusion

By the aid of ColabFold, we succeeded to generate a content-rich structure dataset for reef-building corals within an acceptable period. Its claim to make protein folding accessible to all is seemingly not an exaggeration, furthermore the relatively high confidence of our results might prove that ColabFold does not sacrifice too much accuracy for speed [21].

Our work is expected to be an early case of digital structural proteome building, not just a contribution to coral research. Computational biologists preferring similar approaches may optimize their resource allocation referred to our experience. Moreover, COVID-19 could threat our lives and hamper our fieldwork, but it can’t kill our inspiration or block our communication. Moving most preliminary work to “dry lab” will not only accelebrate research progress via HPC technology, but also help to break regional restrictions and boost brainstorming through the internet, so as to facilitate problem-solving in a more efficient and economical way. In view that the open source of AlphaFold2 and RoseTTAFold has given birth to ColabFold and brought the dawn of the high-throughput era of structural proteomics, we recommend researchers, especially those interested in unfamous yet important organisms, to build and publish digital structural proteomes, which is conducive to new solutions and questions from a structural perspective, people’s deeper understanding of biodiversity, and attraction of joint efforts.

In addition, the dawn of high-throughput structural proteomics might offer bioinformaticians chances to shine. First, there is still much room for improvement in protein structure modelling algorithms. For example, the maximum size of structure predictions is determined by GPU memory currently. Beyond waiting for the update of GPU devices, bioinformaticians can develop algorithms consuming less computing power or supporting distributed computing, further reducing hardware requirements. Second, the development of high-accuracy structure prediction tools may trigger “data explosion” as happened with the emergence of NGS (next-generation sequencing) technology. It gives a variety of possibilities for downstream analysis as well as data mining, nearly a virgin land for method and software development. Last but not least, web apps to visualize and manage structures will receive more attention, adding beauty and convenience to this era. To summarize, let us perform our duties and embrace the spring of structural proteomics.

## Declarations

### Ethics approval and consent to participate

All coral samples were collected and processed in accordance with local laws for invertebrate protection.

### Consent for publication

Not applicable.

### Availability of data and materials

CP-8382 dataset including all sequences, annotations and structure predictions is available at https://doi.org/10.6084/m9.figshare.20128265.v1, and we sincerely recommend acquiring data via our online database introduced in this paper (http://corals.bmeonline.cn/prot/).

### Authors’ contributions

YZ: experiment, database, writing and editing. TH: data uploading. ZL & JC: reviewing. CH: supervision. XL: project approval. All authors contributed to the article and approved the submitted version.

## Acknowledgements

We are grateful for Dr. Xiaojun Xia’s technical support in our production environment.

We thank the Big Data Computing Center of Southeast University for providing the facility support on the numerical calculations in this paper.

## References

1. Odum HT, Odum EP. Trophic structure and productivity of a windward coral reef community on Eniwetok Atoll. Ecological Monographs. 1955;25(3):291–320. doi:10.2307/1943285

2. Yu KF. Coral reefs in the South China Sea: Their response to and records on past environmental changes. Science China Earth Sciences. 2012;55(8):1217–1229. doi:10.1007/s11430-012-4449-5

3. Moberg F, Folke C. Ecological goods and services of coral reef ecosystems. Ecological Economics. 1999;29(2):215–233. doi:10.1016/S0921-8009(99)00009-9

4. Wilson SK, Graham NA, Pratchett MS, Jones GP, Polunin NV. Multiple disturbances and the global degradation of coral reefs: are reef fishes at risk or resilient? GLOBAL CHANGE BIOLOGY. 2006;12(11):2220–2234. doi: 10.1111/j.1365-2486.2006.01252.x

5. Nakamura M, Okaji K, Higa Y, Yamakawa E, Mitarai S. Spatial and temporal population dynamics of the crown-of-thorns starfish, Acanthaster planci, over a 24-year period along the central west coast of Okinawa Island, Japan. Marine Biology. 2014;161(11):2521–2530. doi:10.1007/s00227-014-2524-5

6. Reimer JD, Kise H, Wee HB, Lee C-L, Soong K. Crown-of-thorns starfish outbreak at oceanic Dongsha Atoll in the northern South China Sea. Marine Biodiversity. 2019;49(6):2495–2497. doi:10.1007/s12526-019-01021-2

7. Magel Jennifer M. T., Dimoff Sean A., Baum Julia K. Direct and Indirect Effects of Climate Change-Amplified Pulse Heat Stress Events on Coral Reef Fish Communities. Bulletin of the Ecological Society of America. 2020;101(3):1–6. Accessed February 9, 2022. doi: 10.1002/bes2.1706

8. Dishon G, Grossowicz M, Krom M, Guy G, Gruber DF, Tchernov D. Evolutionary Traits that Enable Scleractinian Corals to Survive Mass Extinction Events. Sci Rep. 2020;10(1):3903. Published 2020 Mar 3. doi:10.1038/s41598-020-60605-2

9. Guo Z, Liao X, Chen JY, He C, Lu Z. Binding Pattern Reconstructions of FGF-FGFR Budding-Inducing Signaling in Reef-Building Corals. Front Physiol. 2022;12:759370. Published 2022 Jan 4. doi:10.3389/fphys.2021.759370

10. Zhu Y, Liao X, Han T, Chen JY, He C, Lu Z. Symbiodiniaceae microRNAs and their targeting sites in coral holobionts: A transcriptomics-based exploration. Genomics. 2022;114(4):110404. doi:10.1016/j.ygeno.2022.110404

11. Ramos-Silva P, Kaandorp J, Huisman L, et al. The skeletal proteome of the coral Acropora millepora: the evolution of calcification by co-option and domain shuffling. Mol Biol Evol. 2013;30(9):2099–2112. doi:10.1093/molbev/mst109

12. Drake JL, Mass T, Haramaty L, Zelzion E, Bhattacharya D, Falkowski PG. Proteomic analysis of skeletal organic matrix from the stony coral Stylophora pistillata. Proc Natl Acad Sci U S A. 2013;110(10):3788–3793. doi:10.1073/pnas.1301419110

13. Conci N, Lehmann M, Vargas S, Wörheide G. Comparative Proteomics of Octocoral and Scleractinian Skeletomes and the Evolution of Coral Calcification. Genome Biol Evol. 2020;12(9):1623–1635. doi:10.1093/gbe/evaa162

14. Peled Y, Drake JL, Malik A, et al. Optimization of skeletal protein preparation for LC-MS/MS sequencing yields additional coral skeletal proteins in Stylophora pistillata. BMC Mater. 2020;2:8. Published 2020 Jul 16. doi:10.1186/s42833-020-00014-x

15. Tunyasuvunakool K, Adler J, Wu Z, et al. Highly accurate protein structure prediction for the human proteome. Nature. 2021;596(7873):590–596. doi:10.1038/s41586-021-03828-1

16. Jumper J, Evans R, Pritzel A, et al. Highly accurate protein structure prediction with AlphaFold. Nature. 2021;596(7873):583–589. doi:10.1038/s41586-021-03819-2

17. Yee A, Pardee K, Christendat D, Savchenko A, Edwards AM, Arrowsmith CH. Structural proteomics: toward high-throughput structural biology as a tool in functional genomics. Acc Chem Res. 2003;36(3):183–189. doi:10.1021/ar010126g

18. Baek M, DiMaio F, Anishchenko I, et al. Accurate prediction of protein structures and interactions using a three-track neural network. Science. 2021;373(6557):871–876. doi:10.1126/science.abj8754

19. Zhu Y, Lu N, Chen JY, He C, Huang Z, Lu Z. Deep whole-genome resequencing sheds light on the distribution and effect of amphioxus SNPs. BMC Genom Data. 2022;23(1):26. Published 2022 Apr 8. doi:10.1186/s12863-022-01038-w

20. Zhong B, Su X, Wen M, Zuo S, Hong L, Lin J. ParaFold: Paralleling AlphaFold for Large-Scale Predictions. International Conference on High Performance Computing in Asia-Pacific Region Workshops. January 2022:1–9. doi:10.1145/3503470.3503471

21. Mirdita M, Schütze K, Moriwaki Y, Heo L, Ovchinnikov S, Steinegger M. ColabFold: making protein folding accessible to all. Nat Methods. 2022;19(6):679–682. doi:10.1038/s41592-022-01488-1

22. Mirdita M, Steinegger M, Söding J. MMseqs2 desktop and local web server app for fast, interactive sequence searches. Bioinformatics. 2019;35(16):2856–2858. doi:10.1093/bioinformatics/bty1057

23. David A, Islam S, Tankhilevich E, Sternberg MJE. The AlphaFold Database of Protein Structures: A Biologist’s Guide. J Mol Biol. 2022;434(2):167336. doi:10.1016/j.jmb.2021.167336

24. Orsburn BC. Proteome Discoverer-A Community Enhanced Data Processing Suite for Protein Informatics. Proteomes. 2021;9(1):15. Published 2021 Mar 23. doi:10.3390/proteomes9010015

25. Priyam A, Woodcroft BJ, Rai V, et al. Sequenceserver: A Modern Graphical User Interface for Custom BLAST Databases. Mol Biol Evol. 2019;36(12):2922–2924. doi:10.1093/molbev/msz185

26. Sehnal D, Bittrich S, Deshpande M, et al. Mol* Viewer: modern web app for 3D visualization and analysis of large biomolecular structures. Nucleic Acids Res. 2021;49(W1):W431–W437. doi:10.1093/nar/gkab314

27. Eighteen Coral Genomes Reveal the Evolutionary Origin of Acropora Strategies to Accommodate Environmental Changes

28. Nelson MR, Thulin E, Fagan PA, Forsén S, Chazin WJ. The EF-hand domain: a globally cooperative structural unit. Protein Sci. 2002;11(2):198–205. doi:10.1110/ps.33302

29. Wang X, Zoccola D, Liew YJ, et al. The Evolution of Calcification in Reef-Building Corals. Mol Biol Evol. 2021;38(9):3543–3555. doi:10.1093/molbev/msab103

30. Von Euw S, Zhang Q, Manichev V, et al. Biological control of aragonite formation in stony corals. Science. 2017;356(6341):933–938. doi:10.1126/science.aam6371

31. Wheeler AP, George JW, Evans CA. Control of calcium carbonate nucleation and crystal growth by soluble matrx of oyster shell. Science. 1981;212(4501):1397–1398. doi:10.1126/science.212.4501.1397

32. Addadi L, Moradian J, Shay E, Maroudas NG, Weiner S. A chemical model for the cooperation of sulfates and carboxylates in calcite crystal nucleation: Relevance to biomineralization. Proc Natl Acad Sci U S A. 1987;84(9):2732–2736. doi:10.1073/pnas.84.9.2732

33. Maurer P, Hohenester E, Engel J. Extracellular calcium-binding proteins. Curr Opin Cell Biol. 1996;8(5):609–617. doi:10.1016/s0955-0674(96)80101-3

34. Fairman JW, Noinaj N, Buchanan SK. The structural biology of β-barrel membrane proteins: a summary of recent reports. Curr Opin Struct Biol. 2011;21(4):523–531. doi:10.1016/j.sbi.2011.05.005

35. Lima C, Disner GR, Falcão MAP, et al. The Natterin Proteins Diversity: A Review on Phylogeny, Structure, and Immune Function. Toxins (Basel). 2021;13(8):538. Published 2021 Jul 31. doi:10.3390/toxins13080538

36. Magalhães GS, Junqueira-de-Azevedo IL, Lopes-Ferreira M, Lorenzini DM, Ho PL, Moura-da-Silva AM. Transcriptome analysis of expressed sequence tags from the venom glands of the fish Thalassophryne nattereri. Biochimie. 2006;88(6):693–699. doi:10.1016/j.biochi.2005.12.008

37. Dal Peraro M, van der Goot FG. Pore-forming toxins: ancient, but never really out of fashion. Nat Rev Microbiol. 2016;14(2):77–92. doi:10.1038/nrmicro.2015.3

38. Greaney AJ, Leppla SH, Moayeri M. Bacterial Exotoxins and the Inflammasome. Front Immunol. 2015;6:570. Published 2015 Nov 10. doi:10.3389/fimmu.2015.00570

39. Ponting CP, Mott R, Bork P, Copley RR. Novel protein domains and repeats in Drosophila melanogaster: insights into structure, function, and evolution. Genome Res. 2001;11(12):1996–2008. doi:10.1101/gr.198701

40. Unno H, Matsuyama K, Tsuji Y, et al. Identification, Characterization, and X-ray Crystallographic Analysis of a Novel Type of Mannose-Specific Lectin CGL1 from the Pacific Oyster Crassostrea gigas. Sci Rep. 2016;6:29135. Published 2016 Jul 5. doi:10.1038/srep29135

41. Jiang S, Wang L, Huang M, et al. DM9 Domain Containing Protein Functions As a Pattern Recognition Receptor with Broad Microbial Recognition Spectrum. Front Immunol. 2017;8:1607. Published 2017 Nov 29. doi:10.3389/fimmu.2017.01607

42. Wang W, Song X, Wang L, Song L. Pathogen-Derived Carbohydrate Recognition in Molluscs Immune Defense. Int J Mol Sci. 2018;19(3):721. Published 2018 Mar 3. doi:10.3390/ijms19030721

43. Li Y, Orange JS. Degranulation enhances presynaptic membrane packing, which protects NK cells from perforin-mediated autolysis. PLoS Biol. 2021;19(8):e3001328. Published 2021 Aug 3. doi:10.1371/journal.pbio.3001328

44. Rosset SL, Oakley CA, Ferrier-Pagès C, Suggett DJ, Weis VM, Davy SK. The Molecular Language of the Cnidarian-Dinoflagellate Symbiosis. Trends Microbiol. 2021;29(4):320–333. doi:10.1016/j.tim.2020.08.005

45. Davy SK, Allemand D, Weis VM. Cell biology of cnidarian-dinoflagellate symbiosis. Microbiol Mol Biol Rev. 2012;76(2):229–261. doi:10.1128/MMBR.05014-11

46. Weis VM. Cell Biology of Coral Symbiosis: Foundational Study Can Inform Solutions to the Coral Reef Crisis. Integr Comp Biol. 2019;59(4):845–855. doi:10.1093/icb/icz067

